# The landscape of gene loss and missense variation across the mammalian tree informs on gene essentiality

**DOI:** 10.1101/2024.05.16.594531

**Authors:** Calwing Liao, Robert Ye, Franjo Ivankovic, Jack M. Fu, Raymond Walters, Chelsea Lowther, Elise Walkanas, Claire Churchhouse, Kaitlin E. Samocha, Kerstin Lindblad-Toh, Elinor Karlsson, Michael Hiller, Michael E. Talkowski, Benjamin M. Neale

## Abstract

**Background:** The degree of gene and sequence preservation across species provides valuable insights into the relative necessity of genes from the perspective of natural selection. Here, we developed novel interspecies metrics across 462 mammalian species, GISMO (Gene identity score of mammalian orthologs) and GISMO-mis (GISMO-missense), to quantify gene loss traversing millions of years of evolution. GISMO is a measure of gene loss across mammals weighed by evolutionary distance relative to humans, whereas GISMO-mis quantifies the ratio of missense to synonymous variants across mammalian species for a given gene.

**Rationale:** Despite large sample sizes, current human constraint metrics are still not well calibrated for short genes. Traversing over 100 million years of evolution across hundreds of mammals can identify the most essential genes and improve gene-disease association. Beyond human genetics, these metrics provide measures of gene constraint to further enable mammalian genetics research.

**Results:** Our analyses showed that both metrics are strongly correlated with measures of human gene constraint for loss-of-function, missense, and copy number dosage derived from upwards of a million human samples, which highlight the power of interspecies constraint. Importantly, neither GISMO nor GISMO-mis are strongly correlated with coding sequence length. Therefore both metrics can identify novel constrained genes that were too small for existing human constraint metrics to capture. We also found that GISMO scores capture rare variant association signals across a range of phenotypes associated with decreased fecundity, such as schizophrenia, autism, and neurodevelopmental disorders. Moreover, common variant heritability of disease traits are highly enriched in the most constrained deciles of both metrics, further underscoring the biological relevance of these metrics in identifying functionally important genes. We further showed that both scores have the lowest duplication and deletion rate in the most constrained deciles for copy number variants in the UK Biobank, suggesting that it may be an important metric for dosage sensitivity. We additionally demonstrate that GISMO can improve prioritization of recessive disorder genes and captures homozygous selection.

**Conclusions:** Overall, we demonstrate that the most constrained genes for gene loss and missense variation capture the largest fraction of heritability, GISMO can help prioritize recessive disorder genes, and identify the most conserved genes across the mammalian tree.

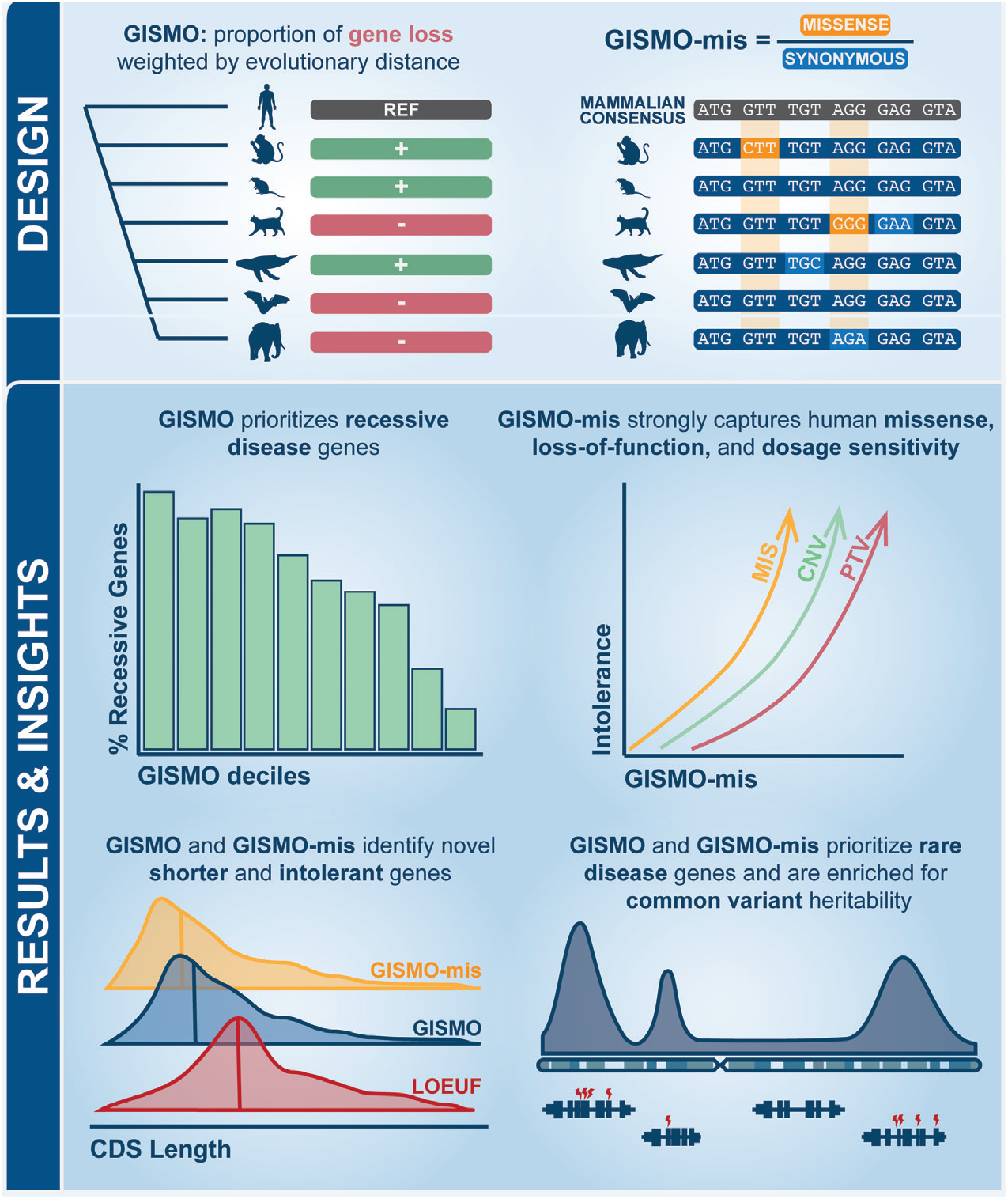

## INTRODUCTION

The application of sequencing technologies at scale has enabled the quantification of gene essentiality in humans, particularly for loss-of-function and missense genetic variants^1,2^. From an evolutionary perspective, not all genes are equally impacted by loss-of-function variants, and some genes are under stronger constraint against such variants than others. Genes that are essential for survival, development and reproduction tend to be more constrained, as damaging and loss-of-function variants in these genes lower fitness and often have severe detrimental effects^3,4^.

Measuring the selective constraint in humans has been made possible by large-scale genomic datasets such as the Genome Aggregation Database (gnomAD)^2^. The gnomAD consortium provides catalogs of discovered genetic variants across >800,000 human individuals, and metrics such as the loss-of-function observed/expected upper bound fraction (LOEUF) scores and missense equivalent (MOEUF) use estimates of the mutation rate alongside patterns of synonymous mutations to calibrate the expected number of loss-of-function and missense mutations observed in the dataset. Loss-of-function and missense mutations that impact genes with important functions will tend not to propagate across generations, leading to fewer such mutations than expected given the mutation rate^5^.

The use of human constraint metrics has already provided valuable insights into the genetic basis of diseases and has been transformative for clinical genetics^6,7^. Genes associated with human genetic disorders that reduce fecundity, such as developmental disorders, are often found to have low LOEUF scores, indicating that these genes carry substantial consequences from a natural selection perspective^8–11^.

One of the main limitations of LOEUF is that genes with shorter sequences have fewer potential loss-of-function mutation sites and attempts to quantify the reduction in such mutations is frustrated by the limited number of potential mutations (i.e., it is difficult to determine whether the number of sites observed is low when the expectation for the number of sites is low itself)^2^. Furthermore, quantifying intraspecies genetic variation requires vastly large sample sizes and can be costly^*2*,12^. One potential solution is to quantify interspecies constraint across the evolutionary tree.

Orthologs are genes that have evolved from a common ancestral gene through speciation events, typically resulting in similar gene sequences and functions across different species^13^. The identification and characterization of orthologous genes has therefore long had an influential role in understanding the evolutionary relationships and functional conservation of genes across different organisms. This is particularly true in mammals where the study of orthologs has enabled the investigation of the genetic basis of various biological processes and diseases^14^. Mammals are a diverse group of organisms that exhibit a wide range of physiological and anatomical characteristics^15,16^. Despite this diversity, mammals share a common ancestry, and studying the orthologous genes across different mammalian species provides insights into the evolutionary changes that have shaped mammalian biology^17^. Comparative genomics studies have shown that orthologs often retain similar functions, suggesting a strong selective pressure to maintain their essential roles throughout evolution^13,18,19^.

Here, we characterized the landscape of gene loss and amino acid (missense) changes across placental mammalian reference genomes, and developed novel metrics for measuring gene loss constraint across 462 mammalian species that cover ∼10% of all recognized species in this clade: GISMO (Gene identity score of mammalian orthologs). We additionally developed GISMO-mis (GISMO-missense), which captures the missense to synonymous ratio of a given gene. We find that both metrics capture signals from existing human constraint metrics such as LOEUF/MOEUF, as well as measures of dosage sensitivity for deletion and duplication. GISMO works across longer timescales and therefore has the ability to address issues with short genes in intraspecies metrics such as LOEUF. can help identify shorter constrained genes. We further demonstrate that GISMO can capture common variant heritability and rare variant association to traits with lower fecundity such as neurodevelopmental disorders (NDDs). Finally, we highlight how GISMO can prioritize recessive disease genes and taps into both heterozygous and homozygous selection. Overall, our findings will contribute to our understanding of the intrinsic properties of genes, the maintenance of essential gene functions, and gene prioritization for human disease.

## RESULTS

### Characterization of gene loss, missense and synonymous variation across mammals

To characterize the distributions of gene loss, missense and synonymous variation, we initially analyzed a comprehensive ortholog data set compiled for 462 placental mammalian species. Since we aim at better understanding human gene constraint, we used the human gene annotation as a reference and the TOGA method to infer orthologs in other mammals^20^. TOGA also provides codon alignments, which we used to detect synonymous and missense mutations, and detects gene-inactivating mutations (frameshift, stop codon, splice site mutations and larger deletions) that we use to detect potential gene loss events in other mammals. We use the term gene loss throughout the manuscript to indicate the absence of a gene encoding an intact reading frame in other mammals, even though our dataset includes human- or primate-specific genes that never existed outside of primates.

For example, 12 genes have been lost across all species except humans, which were typically pseudogenes or single exon genes (**Supplementary Table 1**).

We found that each gene had a median of 18 gene loss events (**Figure 1A**), whereas across the species, we find a median of 1130 likely gene loss events per species (**Figure 1D**). Here, we define gene loss as disruption of the open reading frame. We identified only six genes with no instances of gene loss across the 462 mammalian species: nucleosome assembly protein 1 like 1 (*NAP1L1*), zinc finger CCCH-type containing 14 (*ZC3H14*), Meis homeobox 2 (*MEIS2*), striatin interacting protein 1 (*STRIP1*), chaperonin containing TCP1 subunit 5 (*CCT5*), and ATPase H+ transporting V0 subunit d1 (*ATP6V0D1*).

**Fig. 1.**
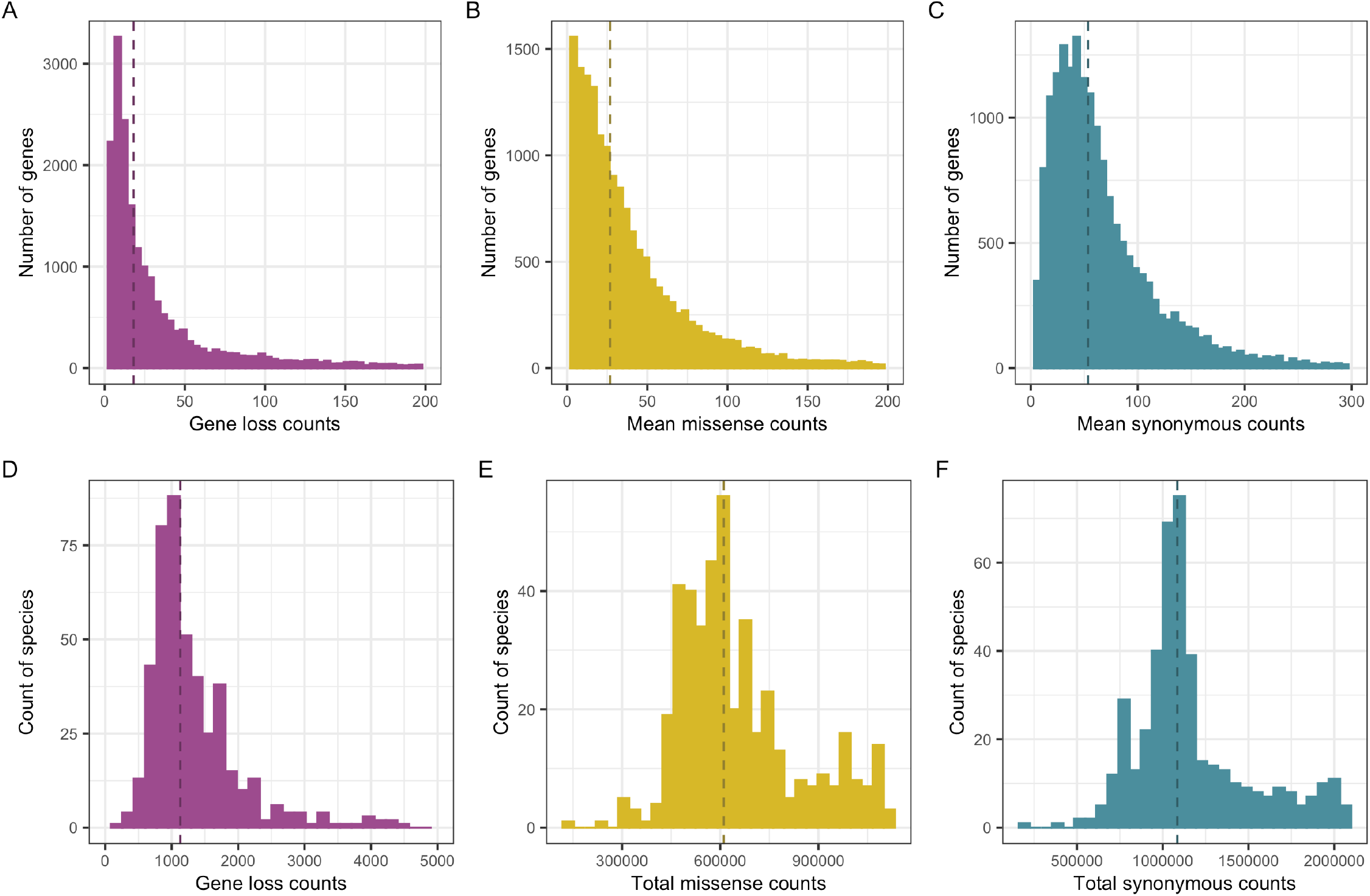
Characterization of gene loss, missense and synonymous counts across 462 placental mammals. (A) Distribution of gene loss. (B) Mean missense variant counts across genes relative to the mammalian consensus. (C) Mean synonymous variant counts across genes relative to the mammalian consensus. (D) Frequency of gene loss per species. (E) Total missense variant counts per species. (F) Total synonymous variant counts per species.

For each gene, we calculated the mean missense and synonymous count across the mammals. The median missense average was 27.7 (**Figure 1B**), whereas the median synonymous average was 53.4 (**Figure 1C**). For each species, there was a median of 610,603 missense variants (**Figure 1E**) and 1,084,983 synonymous variants (**Figure 1F**) across all genes.

### Defining constraint

To capture human gene essentiality, we developed the novel metric, GISMO (Gene Identity Score of Mammalian Orthologs), which estimates the proportion of gene loss that occurs across the 462 mammals surveyed, weighted by evolutionary distance relative to humans. GISMO is a quantitative metric, where a smaller value indicates less frequent gene loss across mammalia (**Supplementary Table 2**). Next, we sought to quantify the missense differences across mammalia relative to a mammalian consensus sequence, a proxy for molecular adaptation and selection of genes. We calculated a second metric, GISMO-mis, which measures the mean missense to synonymous ratio across mammalia (**Supplementary Table 3**).

We evaluated both metrics against other measures of biological and functional importance of genes. The most constrained GISMO and GISMO-mis deciles had the highest enrichment of genes that cause lethality in mouse knockouts and considered essential in cell lines (**Figures 2A-C**). The least constrained deciles have the highest proportion of non-essential and olfactory genes (**Figure 2 D-E**). We also found that genes that were frequently lost across the evolutionary tree were expressed in less tissues across GTEx (**Supplementary Figure 1**).

**Fig. 2.**
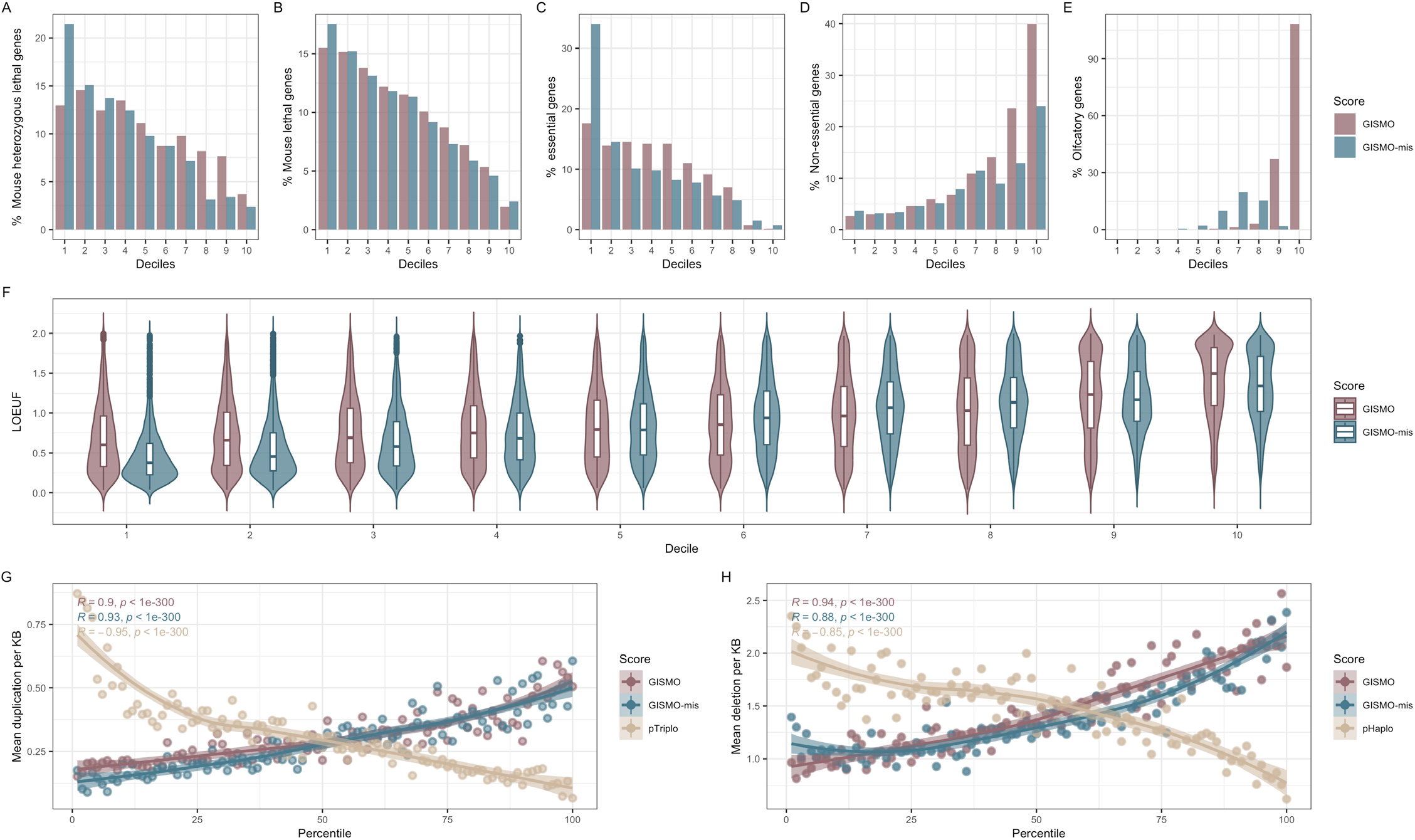
The biological and functional spectrum of gene loss and fixed missense differences. (A-E) The percentage of genes present in each decile of GISMO and GISMO-mis across different genesets (mouse heterozygous lethal, mouse lethal, essential, non-essential, and olfactory genes). The first decile indicates the most constrained. (F) Comparison of GISMO and GISMO-mis to human loss-of-function constraint. (G-H) Comparison of GISMO and GISMO-mis to copy number variants in the UK Biobank (G: duplication, H: deletions). For pHaplo and pTriplo, a higher percentile indicates more constrained whereas for GISMO and GISMO-mis a higher percentile indicates less constrained.

We compared both metrics against human constraint metrics and found that GISMO (R = 0.38, P < 1E-300) and GISMO-mis (R = 0.56, P<1E-300) were both strongly associated with LOEUF (**Figure 2F, Supplementary Figure 2**), with GISMO-mis having a stronger association. Both metrics also had strong correlations with missense constraint (MOEUF), which was comparable to LOEUF for GISMO-mis (R = 0.61, P < 1E-300) but lower for GISMO (R = 0.33, P < 1E-300). We additionally found that both metrics were strongly associated with dosage sensitivity metrics to predict the strength of selection against loss of a gene copy (pHaplo; R GISMO = -0.32., P < 1E-300; R GISMO-mis = -0.44, P < 1E-300) and duplication of a gene (pTriplo; RGISMO = -0.32, P < 1E-300; R GISMO-mis = -0.50, P < 1E-300). We additionally compared GISMO-mis with the new gene-level AlphaMissense metric and found a stronger correlation to this metric (R = 0.84, P < 1E-300) than LOEUF or missense constraint.

Next, to assess the relationship between copy number variants (CNVs) and mammalian gene constraint, we leveraged a CNV dataset generated from exome sequencing in the UK Biobank^21^ (N = 197,306) that enabled detection of individual gene level variants. The most constrained deciles of both GISMO and GISMO-mis had the lowest number of deletions and duplications. We found that GISMO had an even stronger correlation in the UK Biobank for deletions compared to pHaplo, a dosage-sensitivity metric derived from CNVs aggregated from lower-resolution microarray data across ∼1 million human samples^22^ (**Figure 2G-H**).

### Leveraging GISMO to identify shorter intolerant genes

One pitfall of existing human constraint metrics are the biases with gene length. Particularly, shorter genes have a smaller mutational target and are not well calibrated in many human constraint models. The correlation of GISMO and GISMO-mis with coding sequence (CDS) length was -0.12, P = 3.8E-52 for GISMO, -0.12, P = 4.0E-52 for GISMO-mis, and was several magnitudes lower compared to LOEUF (−0.50, P < 1E-300) (**Figure 3A**). Given that LOEUF may not be well calibrated for many short genes, we sought to assess whether GISMO and GISMO-mis can identify shorter intolerant genes. We defined genes to be GISMO and GISMO intolerance if the genes fell within the top 15% most constrained genes (lowest scores). Both GISMO and GISMO-mis identify much shorter intolerant genes (GISMO median CDS = 1502 base pairs for 2897 genes; GISMO-mis median CDS = 1302 base pairs for 2,502 genes) than LOEUF intolerant genes (median CDS = 2267 base pairs for 2,968 genes). We additionally find that genes predicted to be intolerant by GISMO and GISMO-mis but not LOEUF tend to be much shorter compared to genes that are intolerant to LOEUF and one of the GISMO metrics (**Figure 3B**).

**Fig. 3.**
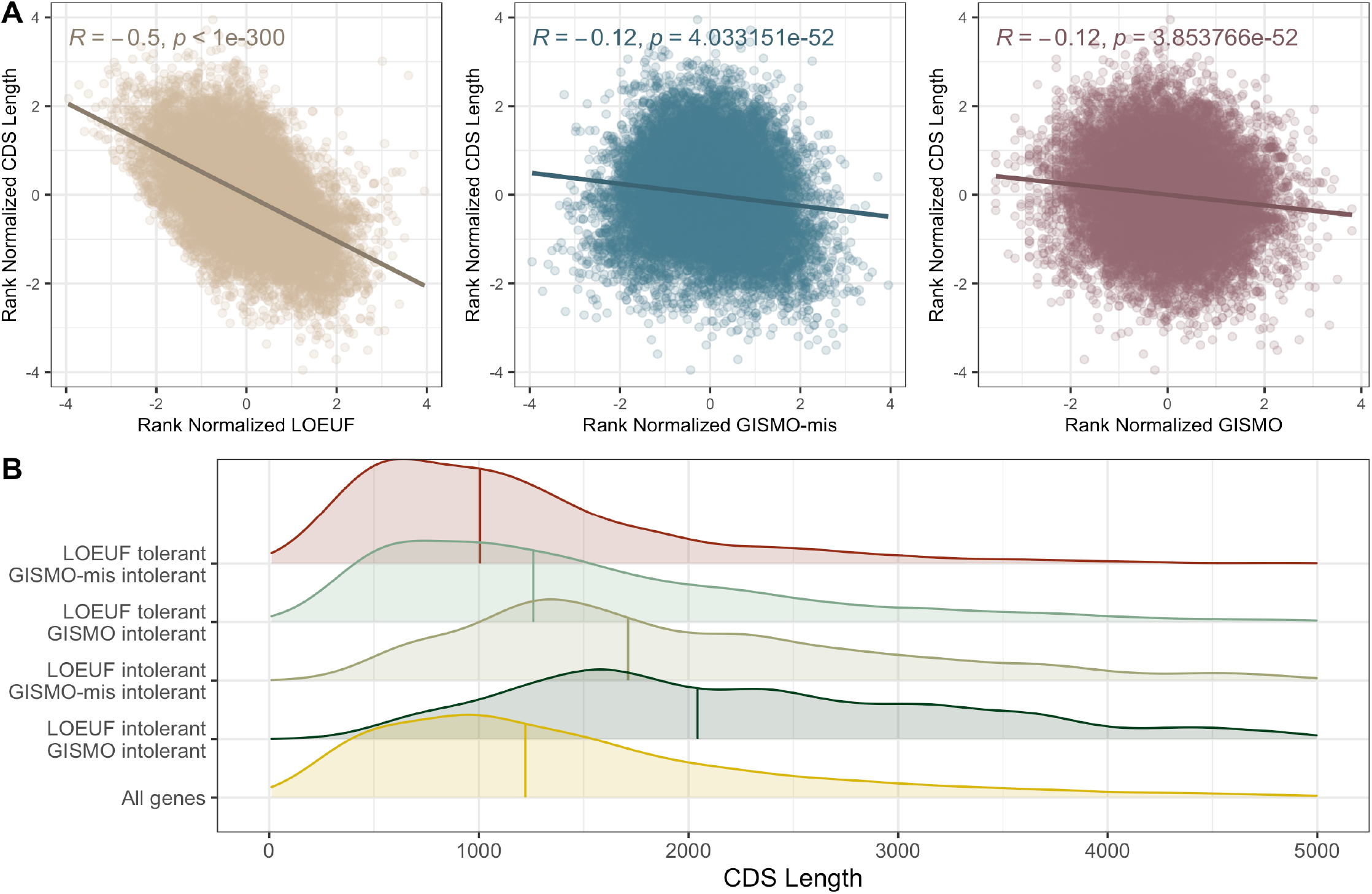
GISMO and GISMO-mis capture shorter genes relative to human constraint metrics. (A) Correlation of GISMO, GISMO-mis and LOEUF against CDS length. The CDS, LOEUF, GISMO and GISMO-mis were inverse rank normalized. A Pearson’s correlation was done. (B) Short constrained GISMO and GISMO-mis genes are considered tolerant for LOEUF. Genes were split into categories of GISMO / GISMO-mis intolerant or LOEUF intolerant. We define intolerance for LOEUF as <0.35 and GISMO/GISMO-mis as the first two most constrained deciles respectively.

The genes that were considered tolerant for LOEUF (>0.35) and GISMO intolerant were enriched for genes involved in the following pathways/ gene sets in mice: sperm immotility (Bonferroni-adjusted P = 2.17E-03, [23/46 genes]), absent acrosome (Bonferroni-adjusted P, 4.86E-2, [16/30 genes]), abnormal actin cytoskeleton morphology (Bonferroni-adjusted P = 4.86E-02, [15/27 genes]) (**Supplementary Table 4**). Intriguingly, when we explored enrichment of these genes amongst human traits, we found phenotypes related to later onset of disease age, which may reflect changes in fecundity and recent selective pressures due to modern medicine, such as abnormal posterior eye segment morphology (Bonferroni-adjusted P = 8.3E-02, [328/1439 genes]) and lipid accumulation in hepatocytes (Bonferroni-adjusted P = 1.0E-01, 50/149).

### Mammalian gene loss and fixed missense differences capture human disease signals

Next, we evaluated whether mammalian gene loss and fixed missense differences could help prioritize genes relevant to human diseases. Given that many disease models are mammals (i.e. mus musculus), assessing whether disease genes are often strongly conserved may help pinpoint what human genes may be reasonably modeled in mammals. First, we assessed common variant heritability. For both metrics, we partitioned heritability across 276 independent traits from the UK Biobank and across several disease traits not well captured in the biobank. We found that there was a strong linear enrichment of heritability across the deciles of both metrics, with lowest enrichment in the least constrained deciles (**Figure 4A**). Second, we looked at rare variant association studies of traits with decreased fecundity, such as autism, schizophrenia, and neurodevelopmental disorders. We found that both GISMO and GISMO-mis constrained genes had a higher association to neurodevelopmental disorders and the metrics were strongly correlated with the association strength (**Figure 4B, Supplementary Figure 3-4**).

**Fig. 4.**
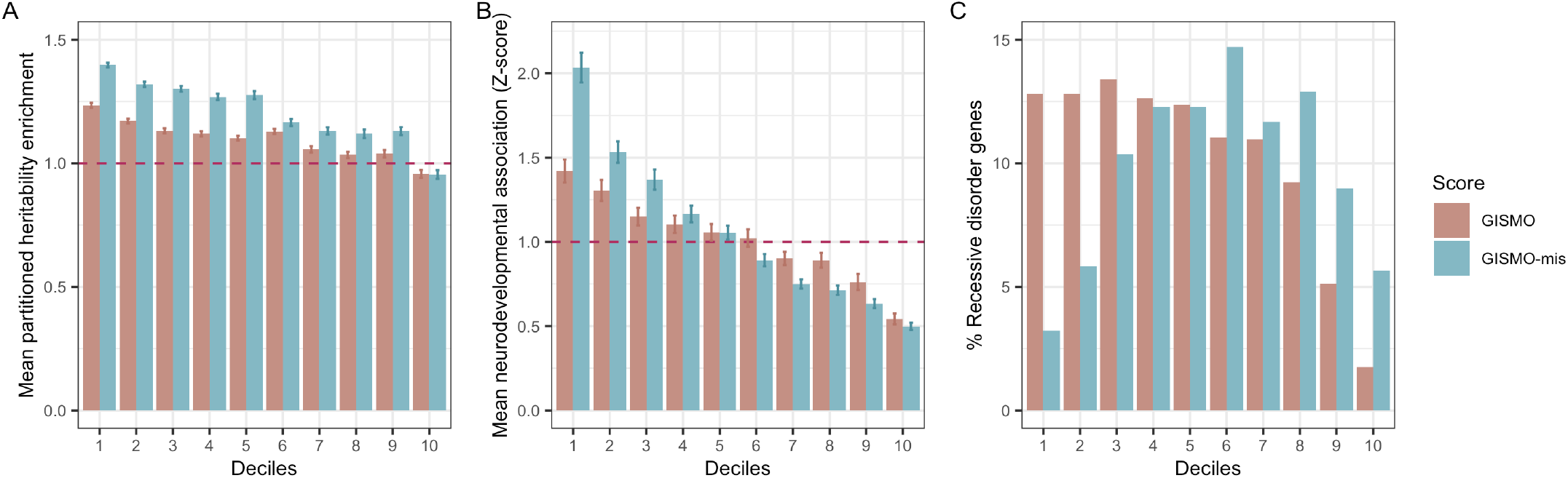
GISMO and GISMO-mis capture common variant heritability and rare variant association signals. (A) Mean enrichment of per-SNV partitioned heritability of XX traits explained by common variants within 100-kb of genes for GISMO and GISMO-mis deciles. (B) Mean neurodevelopmental disorder association across GISMO and GISMO-mis deciles. The association reflects P-values from NDD rare variant associations that were converted to absolute Z-scores. (C) The percentage of genes present in each decile of GISMO and GISMO-mis for 1,183 recessive disorder genes aggregated from OMIM.

### Improving recessive disease gene prioritization

Given that gene loss in a species can be reflective of selection on homozygous loss-of-function carriers, we hypothesized that GISMO could be used to prioritize recessive genes associated to disease (i.e. genes under homozygous selection should be more constrained by GISMO). To assess this, we assembled a list of 1,183 recessive genes and found that there was enrichment amongst the most constrained deciles (**Figure 4C**), whereas GISMO-mis expectedly did not have strong enrichment. Next, we sought to test whether GISMO may help with prioritization of recessive inheritance candidate genes in the Deciphering Developmental Disorder

Study consisting of 14,000 trios. We found that the GISMO distribution for recessive developmental disorder genes with a high confidence (confident or strong) followed similar distributions as Mendelian recessive genes, whereas the recessive genes with limited and moderate evidence were unclear (**Supplementary Figure 5**). We additionally found that the limited and moderate evidence categories prioritized genes with high constraint for LOEUF, a reflection of strong heterozygous selection, which are unlikely gene candidates for recessive disorders.

## DISCUSSION

Evolution and selection are powerful mechanisms that provide insights into essential biology. Comparative genomics leverages millions of years of evolution and selection to understand the importance of genes across various environmental influences. Here, we characterized gene loss and fixed missense differences across the placental mammalian tree, including genomes that cover ∼10% of all recognized species. We found that as expected, most genes are present in the majority of mammals, yet any given species may have upwards of over a thousand genes lost. This suggests that certain gene loss is concentrated in a highly select few genes that are tolerant to this loss. Similarly, missense fixation events tend to be low for most genes, and expectedly fixation differences occur more frequently in genes that are less important for biology. We developed novel metrics, GISMO and GISMO-mis, based on the current largest and most comprehensive set of mammalian species representing roughly 10% of mammals to quantify these selection measures.

GISMO is the first metric, to our knowledge, to quantify gene loss across the mammalian tree. We found that GISMO is well saturated, given that only 6 genes did not have any gene loss events. Amongst these 6 conserved genes, we found that they were related to DNA replication, polyA binding for RNA, and development. In contrast to human-based metrics from gnomAD such as LOEUF and pLI, where over 1000 genes do not have an observed loss of function event across 141,456 individuals in gnomAD v2.1 and 832 genes across 807,162 individuals in gnomAD v4.0. We found that most genes that were likely to be human-specific tended to be enriched for single exon and pseudogenes, likely reflecting potential noise or events that are not real or biologically meaningful. We benchmarked these metrics against known gene sets and model organism data and highlighted the potential for identifying disease genes and function.

There are several advantages of harnessing mammalian biology in the form of these metrics. We demonstrated that both GISMO and GISMO-mis can identify shorter constrained genes under selection, which are much shorter than LOEUF constrained genes. In biology, gene essentiality does not depend on gene length; new metrics that capture gene essentiality without biases from gene length are quintessential for understanding the biological and functional spectrum. We also show that GISMO is able to help prioritize recessive disorder genes, which most constraint metrics are not well calibrated to do. Prior studies have highlighted how recessive metrics such as pRec, which measures probability of a gene being recessive, cannot differentiate weak selection on heterozygotes from homozygotes, a limitation of human genetic data^23^. We further show that combining GISMO and heterozygous selection metrics such as LOEUF can help improve clinical prioritization of genes. For instance, a gene candidate is much more likely to be recessive if the GISMO score is low and the LOEUF score is high. We additionally find that GISMO intolerant and LOEUF tolerant genes are important for fertility and reproductive success, with pathway enrichments such as immotile sperm in mice.

We further emphasize that the strong correlations between GISMO and human constraint metrics highlight the power of interspecies over intraspecies metrics. Particularly, the high costs and sample sizes required to generate calibrated intraspecies human constraint metrics such as LOEUF (>800,000 individuals) and pHaplo/pTriplo (>1 million individuals) in contrast to 462 mammals for GISMO.

Moreover, in human genetics, constraint metrics have been transformative for many different subfields. In particular, the probability of loss-of-function intolerance (pLI) and subsequent loss-of-functional observed/expected upper bound fraction (LOEUF) metrics, derived from large-scale sequencing studies, have become standard parts of genetic and genomic analysis pipelines. These metrics have revolutionized a number of workflows in human genetics, including filtering and prioritization of genes and how likely they are to have phenotypic impact for association studies, identification of which genes or variants are likely under heterozygous loss-of-function selection, as well as clinical interpretation of patient variation. Importantly, the GISMO metrics will allow non-human mammalian researchers to have constraint metrics to further enable and advance the analytical framework in the large field.

An open question from these analyses was whether mammalian gene loss and fixed missense differences could be leveraged to capture important biological insights into human diseases and traits. To explore this question, we analyzed an array of quantitative traits, human disease phenotypes, and gene sets robustly associated with rare and common diseases. We found significant common variant heritability enrichment across a vast number of traits in the most constrained deciles of both GISMO and GISMO-mis. We similarly find both metrics capture rare variant association to traits with decreased fecundity. We posit that GISMO and GISMO-mis can provide orthogonal levels of disease evidence and may help with increasing disease association power. Moreover, we show that the majority of rare and common variant heritability is concentrated in genes that are strongly conserved across species. This reinforces the potential utility of mammalian models for dissection of heritable quantitative traits in humans.

Despite the increased power and clear value demonstrated herein for derived metrics based on mammalian orthologs such as GISMO and GISMO-mis, there remains several limitations in the derivation of these metrics. Mammalian gene conservation may not reflect the human selective pressure and recent innovation that has affected the selection regime of certain traits. It is also important to understand the practical use of GISMO given it taps into both homozygous and heterozygous selection; pairing with additional constraint metrics such as LOEUF and GISMO-mis can help tease apart these relationships. Moreover, the species included may be subjected to survivorship bias and accessibility bias, so further sampling of a broader spectrum of mammals will improve metrics like GISMO and GISMO-mis.

Overall, we demonstrate that mammalian gene loss and missense fixation are important measures of selection. We developed several powerful metrics to quantify evolutionary constraints for gene loss (GISMO) and molecular adaptation (GISMO-mis). Our estimates provide informative ranking of gene importance, which ultimately allow us to better understand gene essentiality and disease association.

## METHODS

### Development of the GISMO metric

To infer orthologs, the Tool to infer Orthologs from Genome Alignments (TOGA)(20) was used with the human GENCODE 38 annotation^24^ across 462 mammals. Since for some species TOGA data was available for multiple assemblies, we selected only the assembly with the highest contig N50. An orthologous gene is considered lost when and only when all transcripts of the respective gene are categorized as lost. To assess the type of orthology, TOGA subsequently evaluates, for each human reference gene, the classification of all its corresponding orthologous loci and which reference genes were annotated.

GISMO was calculated with the following formula:

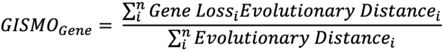

A 95% confidence interval was simulated using a binomial distribution and the upper 95% confidence interval was used. Briefly, the binomial probability was estimated for each phylogenetic order and counts were simulated 10,000 times for the 462 species. Each count was subsequently weighted by the evolutionary distance relative to humans, where 1 = gene loss. Evolutionary distance was standardized by dividing by the maximum evolutionary distance amongst the mammals used to generate GISMO.

### Development of the GISMO-mis

To calculate GISMO-mis, multiple codon alignments including up to 462 mammals have been generated using MACSE v2^25^ and were downloaded from http://genome.senckenberg.de/download/TOGA/. To generate codon alignments, the following selection procedures were performed: 1) Human transcript with the longest coding sequence length, 2) Orthologs were considered if they were classified as intact, partial intact, or uncertain loss. To ensure alignments were mostly 1 to 1 orthologs, if a query species has more than four predicted orthologs, this species was not included in the multiple codon alignment. Additionally, if the gene did not have a single ortholog for at least 75% of all query species, a multiple codon alignment was not computed for this gene. Subsequently for each gene, a mammalian consensus transcript sequence across all species was generated and used as a reference. To determine each consensus transcript sequence, all mammalian sequences for that gene were considered at a codon-level resolution and the most represented codon(s) across all queried species was chosen. In the event of tied codon counts, the consensus sequence permits multiple possible reference codons. Next, missense and synonymous counts across each species were quantified against the mammalian consensus sequence for each transcript. Any codon comparisons involving gaps in either reference or queried codons were not considered. In the case of multiple reference codons at a given position, missense and synonymous counts were conservatively generated by only classifying a queried codon as missense if none of the reference codons were synonymous with the queried codon. Similarly in the case of a codon with ambiguous nucleotides, each ambiguous nucleotide was exhaustively replaced until either 1) a synonymous result was achieved, after which the query would be classified as synonymous, or 2) no synonymous result was achieved across all potential substitutions, after which the query would be classified as missense. Finally for each gene, the mean missense to synonymous ratio across all queried species was calculated.

### Benchmarking against gene sets and pathways

To benchmark GISMO, we compared GISMO against several independent gene sets: lethal mouse, olfactory, essential genes, and non-essential genes. These were the same genesets previously used in gnomAD benchmarking to have a reasonable comparison^2^. Both GISMO and GISMO-mis were split into deciles, where the lowest decile (1st) represented the most constrained. Additionally, data from GTEx 53 was used to assess how expressed genes are across the different number of tissues. Genes were considered expressed in a tissue with a TPM > 0.3. Pathway enrichment was done using https://toppgene.cchmc.org/. A recessive gene set was curated from OMIM^26^, which included a total of 1,183 autosomal recessive genes.

### Correlation between GISMO and GISMO-mis against independent constraint metrics

To assess whether GISMO and GISMO-mis are associated with different constraint scores, both metrics were correlated against other gene-level metrics. First, the GISMO metrics were compared against two human constraint metrics derived from gnomAD for loss-of-function constraint (LOEUF) and missense constraint (MOEUF). Next, to test whether GISMO and GISMO-mis can capture copy number variant dosage sensitivity, the scores were benchmarked against pTriplo (triplosensitivity) and pHaplo (haploinsufficiency)^22^. We additionally benchmarked GISMO and GISMO-mis against gene-level AlphaMissense scores^27^, which measures the effects of missense variation on predicted structural context from AlphaFold. A Spearman’s correlation was used for all correlations.

### Partitioned heritability across independent traits

To assess the distribution of heritability enrichment across GISMO and GISMO-mis, LD score regression (LDSC) was used to partition heritability across gene deciles of both metrics^28–30^. For both metrics, we included a 100kb flanking region both up and downstream for each gene and, in conjunction with genotype data from the 1000 Genomes Project. The SNPs were restricted to HapMap3 SNPs with an estimated annotation-specific LD scores using a 1cM window. Next, partitioned heritability was applied to 276 independent traits from the UK Biobank (https://www.nealelab.is/uk-biobank/), as well as additional disease traits such as schizophrenia, bipolar disorder, autism spectrum disorder, attention-deficit hyperactivity disorder, and coronary artery disease^31–35^. Phenotypes were selected from the UK Biobank based on having a significant p-value (P < 0.05) after Bonferroni correction. Additionally, phenotypes were assessed for phenotypic correlation and only independent traits were included. The baseline model (v2.2), which includes 74 annotations to capture genomic properties, was included alongside the estimated LD scores as a predictor in the LD Score regression. HapMap3 SNPs, excluding the HLA region, were used as default regression SNPs.

### Assess GISMO and GISMO-mis coding sequence biases

To assess whether GISMO and GISMO-mis are biased by coding sequence length, we correlated against the coding sequencing length (CDS) of each gene in both metrics. We compared against the gold standard for loss-of-function constraint, LOEUF from gnomAD. Given that LOEUF had a significantly stronger correlation with CDS relative to GISMO and GISMO-mis, we hypothesized that the GISMO metrics may help prioritize short essential genes. We considered genes with a LOEUF score <0.35 as LOEUF-constrained and intolerant, representing roughly the top 15% most constrained genes. Similarly, for both GISMO and GISMO-mis, we took the top 15% most constrained genes, which we considered intolerant. We compared the CDS length distribution for GISMO and GISMO-mis intolerant and LOEUF tolerant.

## ACKNOWLEDGEMENTS

C.L. is funded by the CIHR Banting Fellowship. K.L.T is a Distinguished professor funded by the Swedish Research Council. M.H was supported by the LOEWE-Centre for Translational Biodiversity Genomics (TBG) funded by the Hessen State Ministry of Higher Education, Research and the Arts (HMWK).

## AUTHOR CONTRIBUTIONS

C.L. and B.M.N. came up with the project. C.L. and R.Y. conducted all analyses and wrote the manuscript. F.I., J.M.F., C.L, E.V. provided analytical and scientific assistance. B.M.N., M.E.T. supervised the project. K.E.S., K.L.T., E.K., M.H., C.C., R.W. provided additional supervision and edits to the manuscript.

## CODE AND DATA AVAILABILITY

All code and relevant data are available on GitHub: https://github.com/cliao5/GISMO/.

