## Supplementary Figures and Text for "The landscape of gene loss and missense variation across the mammalian tree informs on gene essentiality"

**Supplementary Text and Figures**

Development of the GISMO metric

First, we had to infer orthologs across existing mammalian assemblies. A large portion of new mammalian assemblies were derived from Zoonomia. To infer orthologs, the Tool to infer Orthologs from Genome Alignments (TOGA)(*20*) was used with the human GENCODE 38 annotation(*24*) across 462 mammals. Since for some species TOGA data was available for multiple assemblies, we selected only the assembly with the highest contig N50. An orthologous gene is considered lost when and only when all transcripts of the respective gene are categorized as lost. To assess the type of orthology, TOGA subsequently evaluates, for each human reference gene, the classification of all its corresponding orthologous loci and which reference genes were annotated. Ortholog classification results across 462 mammals (<https://genome.senckenberg.de/download/TOGA/human_hg38_reference/>), were used to construct a matrix cataloging every gene from every mammalian species as having a one-to-one, one-to-many, one-to-zero, many-to-one, or many-to-many ortholog correspondence.

 GISMO was calculated with the following formula:

 
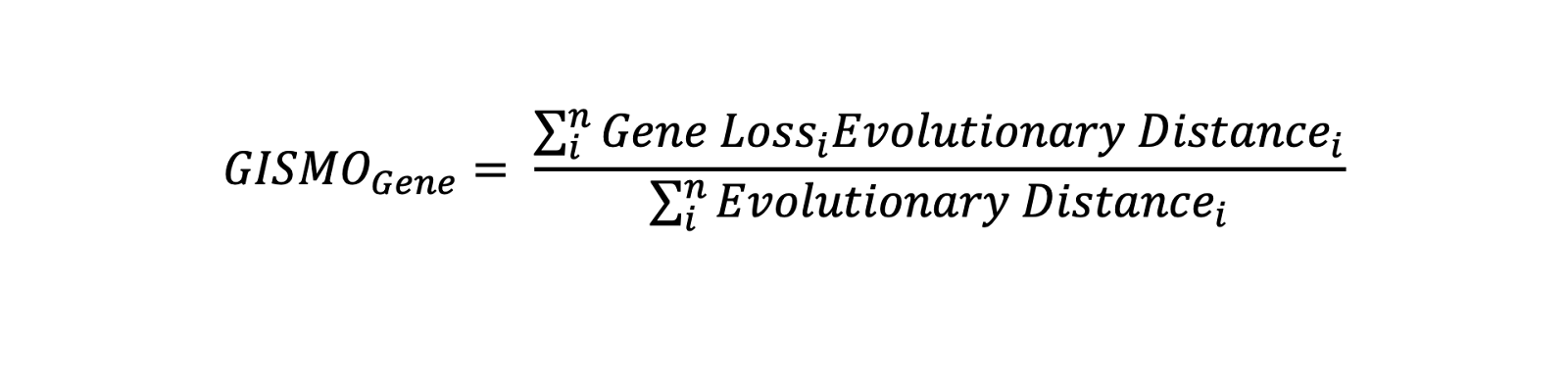

 A 95% confidence interval was simulated using a binomial distribution and the upper 95% confidence interval was used. Briefly, the binomial probability was estimated for each phylogenetic order and counts were simulated 10,000 times for the 462 species. We chose to use the upper confidence bound to make each gene score more conservative. Each count was subsequently weighted by the evolutionary distance relative to humans, where 1 = gene loss. Evolutionary distance relative to humans was standardized by dividing by the maximum evolutionary distance amongst the mammals used to generate GISMO. We compared the metric with and without evolutionary distance weighting. We decided to move forward with the evolutionary distance weighting because there were marginal increases in correlation with human constraint metrics.

Benchmarking against gene sets and pathways

To test the robustness of our new scores, we sought to identify independent datasets to determine whether the scores sufficiently capture these independent metrics and expected gene sets. To benchmark GISMO, we compared GISMO against several independent gene sets: lethal mouse, olfactory, essential genes, and non-essential genes. These were the same genesets previously used in gnomAD benchmarking to have a reasonable comparison(*2*). Both GISMO and GISMO-mis were split into deciles, where the lowest decile (1st) represented the most constrained.  Additionally, data from GTEx 53 was used to assess how expressed genes are across the different number of tissues. Genes were considered expressed in a tissue with a TPM >0.3. Pathway enrichment was done using <https://toppgene.cchmc.org/>. A recessive gene set was curated from OMIM(*26*), which included a total of 1,183 autosomal recessive genes.

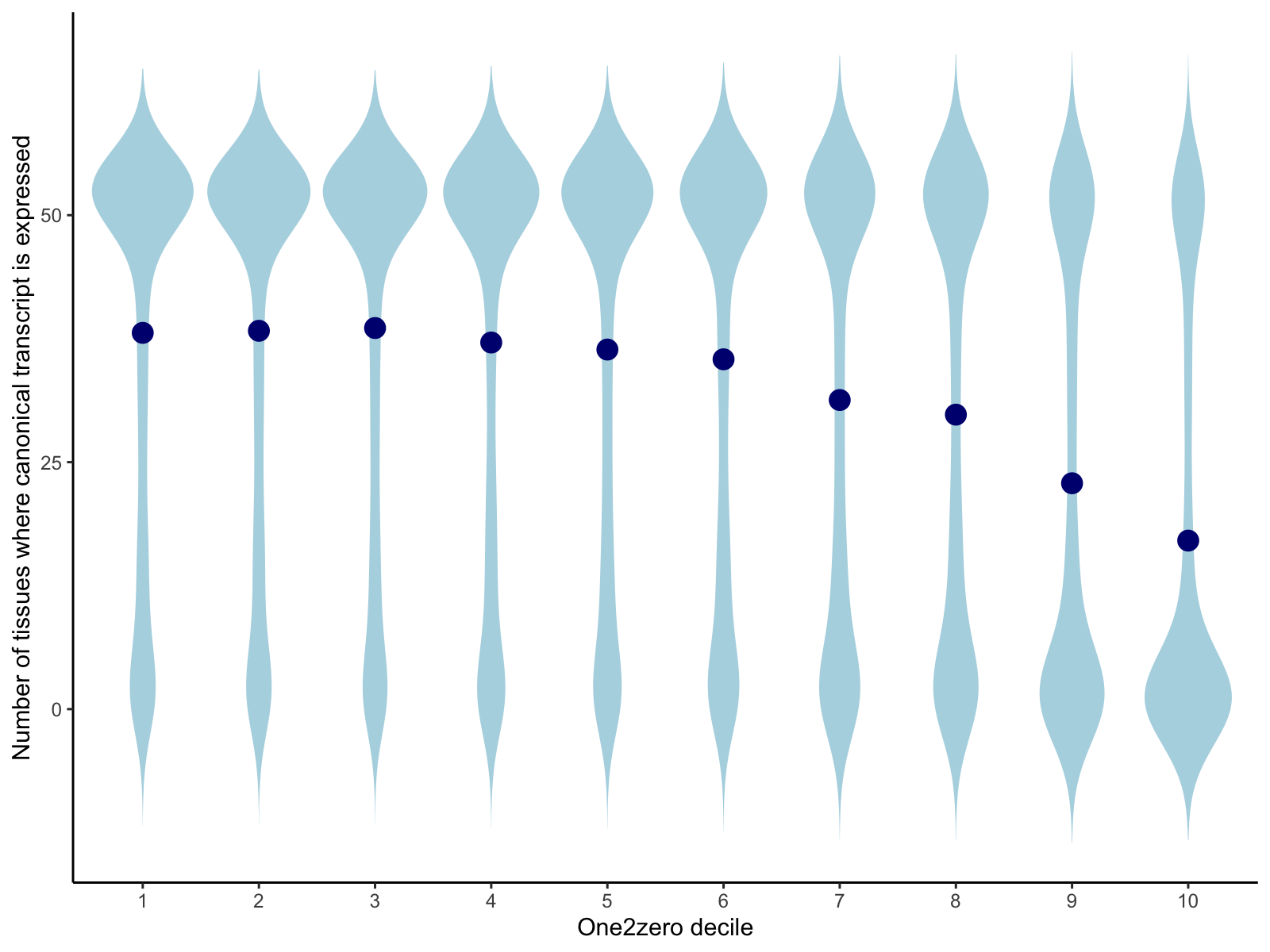

**Supplementary Figure 1. The most frequently lost genes across the mammalian tree tend to be expressed in the least number of tissues across GTEx 53.** The blue point represents the number of tissues that have any meaningful expression (>0.3 TPM) across 53 GTEx human tissues. One2zero is analogous to GISMO and represents a gene loss event, where a higher decile indicates highest gene loss.

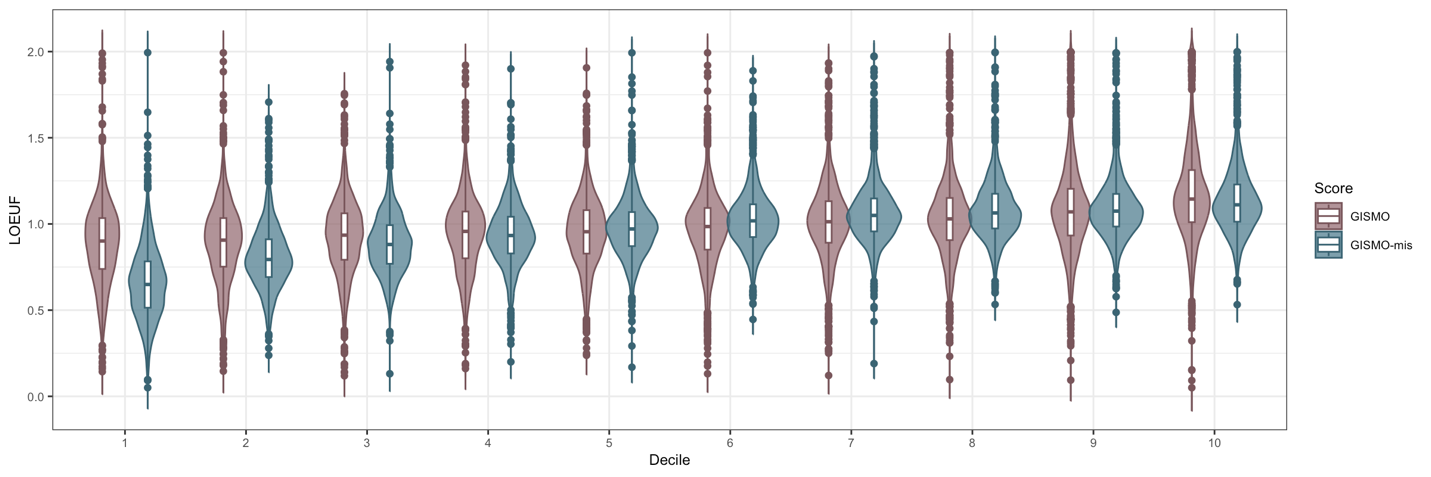

**Supplementary Figure 2. Comparison of GISMO and GISMO-mis to human missense constraint.** Missense constraint is defined as MOEUF from gnomAD, where lower deciles indicate most constrained.

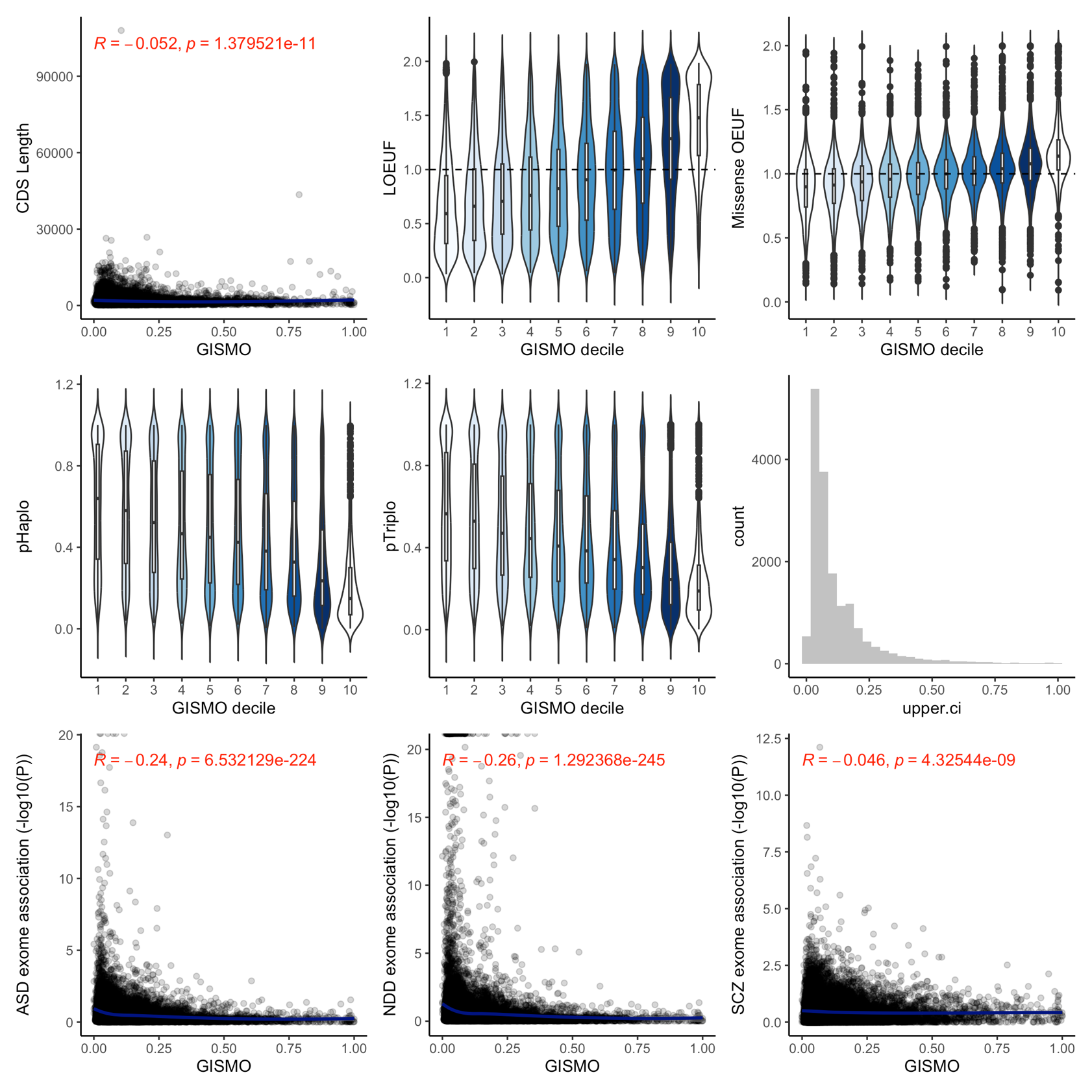

**Supplementary Figure 3. Association between neuropsychiatric rare variant associated genes and GISMO.** A Spearman’s correlation was used.

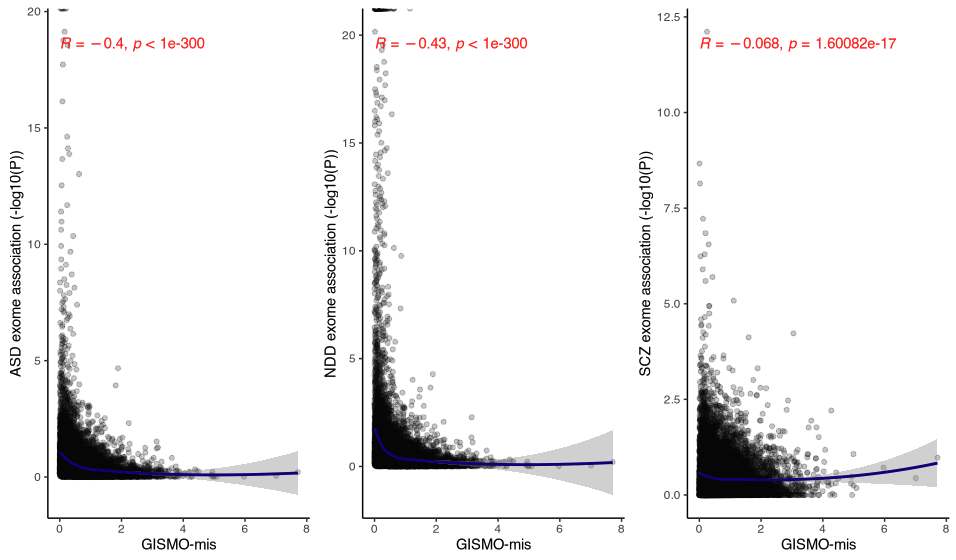

**Supplementary Figure 4. Association between neuropsychiatric rare variant associated genes and GISMO-mis.** A Spearman’s correlation was used.

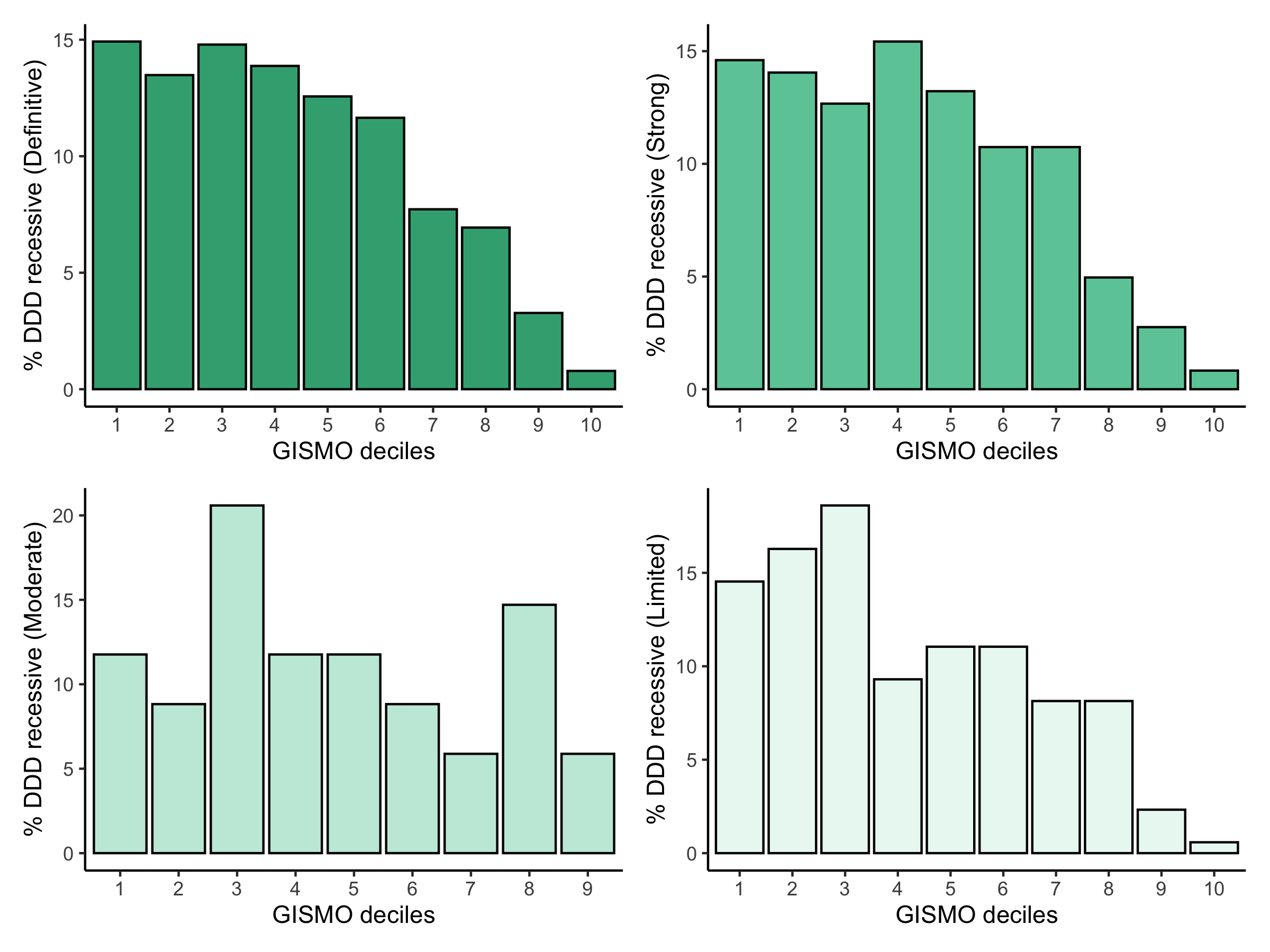

**Supplementary Figure 5. GISMO can help prioritize recessive disorder associated genes from the Deciphering Developmental Disorders (DDD) cohort.**
